# Landscape-scale exposure to multiazole-resistant *Aspergillus fumigatus* bioaerosols

**DOI:** 10.1101/2022.11.07.515445

**Authors:** Jennifer M. G. Shelton, Johanna Rhodes, Christopher B. Uzzell, Samuel Hemmings, Amelie P. Brackin, Thomas R. Sewell, Asmaa Alghamdi, Paul S. Dyer, Mark Fraser, Andrew M. Borman, Elizabeth M. Johnson, Frédéric B. Piel, Andrew C. Singer, Matthew C. Fisher

**Affiliations:** MRC Centre for Global Infectious Disease Analysis, Department of Infectious Disease Epidemiology, Imperial College London, London, UK; UK Centre for Ecology & Hydrology, Wallingford, Oxfordshire, UK; Department of Medical Microbiology, Radboud University Medical Center, Nijmegen, The Netherlands; School of Life Sciences, University of Nottingham, Nottingham, UK; UK National Mycology Reference Laboratory, National Infections Service, Public Health England, Science Quarter, Southmead Hospital, Bristol, UK; MRC Centre for Medical Mycology, University of Exeter, UK; NIHR HPRU in Environmental Exposures & Health, Department of Epidemiology and Biostatistics, Imperial College London, London, UK

## Abstract

We demonstrate country-wide exposures to aerosolized spores of a human fungal pathogen, *Aspergillus fumigatus*, that has acquired resistance to first line azole clinical antifungal drugs. Assisted by a network of citizen scientists across the United Kingdom, we show that 1 in 20 viable aerosolized spores of this mold are resistant to the agricultural fungicide tebuconazole and 1 in 140 spores are resistant to the four most used azoles for treating clinical aspergillosis infections. Season and proximity to industrial composters were associated with growth of *A. fumigatus* from air samples, but not with the presence of azole resistance, and hotspots were not stable between sampling periods suggesting a high degree of atmospheric mixing. Genomic analysis shows no distinction between those resistant genotypes found in the environment and in patients, indicating that ~40% (58/150 sequenced genomes) of azole-resistant *A. fumigatus* infections are acquired from environmental exposures. Due to the ubiquity of this measured exposure, it is crucial that we determine source(s) of azole-resistant *A. fumigatus*, who is at greatest risk of exposure and how to mitigate these exposures, in order to minimize treatment failure in patients with aspergillosis.

**One sentence summary:** UK-wide citizen science surveillance finds a ubiquitous exposure to aerosolized spores of a human fungal pathogen that have evolved in the environment cross-resistance to essential clinical antifungal drugs

## Manuscript

The cosmopolitan mold, *Aspergillus fumigatus*, causes a spectrum of chronic and acute life-threatening diseases in humans. The widespread occurrence of resistance to first line clinical azole drugs in environmental isolates, alongside molecular epidemiology linking azole-resistant *A. fumigatus* (*ARAf*) sourced from patients and the environment, argues that a substantial burden of treatment failure is due to environmental azole exposure (*1*). The numbers of patients in the United Kingdom (UK) presenting with infections that are resistant to one or more of the clinical azoles is increasing in diverse patient groups (*2, 3*). The significantly elevated case fatality rates where invasive aspergillosis is caused by AR*Af* (*4, 5*) further highlights the importance and breadth of this emerging problem.

The widespread use of broad-spectrum agricultural fungicides, founded on the same demethylase inhibitor (DMI) chemistry as the clinical azoles, has long been argued to drive the evolution of environmental resistance (*6*). This hypothesis has found broad support from surveillance demonstrating environmental hotspots of *A. fumigatus* growth alongside high frequencies of resistance where this saprotrophic mold has the potential to grow in the presence of agricultural DMIs (*7*). Yet, despite its increasingly wide detection in the environment worldwide (*7, 8*), little is known about the extent to which humans are exposed to AR*Af*. The mold is adapted to airborne dispersal and most humans inhale large numbers of viable spores every day (*9, 10*). Owing to the potential clinical consequences of their inhalation, occupational exposures to these spores are legislated in countries such as the UK, especially in green-waste recycling and composting processes that have the potential to generate high levels of inocula (*11*). However, occupational monitoring does not include assessing exposures to AR*Af*.

Surveillance has shown hotspots of environmental resistance in environments including both home (*8*) and industrial compost (*12*), urban environments (*13*), greenhouses (*14*) and horticultural products (*15*). Nonetheless, there is little insight into population-wide exposures of at-risk individuals to aerosolised AR*Af* occurring beyond these heterogeneous environmental foci.

Given the dynamic nature of the atmosphere and the potential for season to affect the biology of *A. fumigatus* spore production, meaningful assessment of human exposure to AR*Af* needs to be undertaken at a population level in a cost-effective manner that spans yearly seasonal variation. To meet this need, a UK-wide campaign was launched using social media, and used to recruit a network of citizen scientists (*16*). These individuals then used simple passive air samplers to collect airborne spores of *A. fumigatus* synchronously across a 6-10-hour time period on the days matching the northern hemisphere seasonal equinoxes and solstices between 2018 and 2019 (**Figure 1A–C**). This activity resulted in a total of 1,894 air samples being collected that, whilst being clustered owing to greater sample collection in areas of high population density, achieved a near UK-wide distribution at each timepoint (**Figure 2A–D**). These air samples were incubated on growth media using the highly selective temperature of 43°C and 919 (49%) yielded a combined 2,366 *Aspergillus* colonies. Of these, secondary screening on media containing 6 mg/L of the commonly-used agricultural fungicide tebuconazole (TEB; (*17*)) identified 111 TEB-resistant isolates, comprising 4.7 % of the total isolates recovered (**Table 1**). This frequency is similar to that (~4%) measured at the Rothamsted Research station in 2016 suggesting an incidence that is relatively stable across recent years (*18*). Of the TEB-resistant isolates, 12 failed to sequence using the *cyp51A* promoter and coding region primers and were re-identified by MALDI-TOF mass spectrometry as the related species *Aspergillus lentulus* (*n* = 10) and *Aspergillus nidulans* (*n* = 2). For the 99 TEB-resistant isolates confirmed to be *A. fumigatus*, clinical breakpoints then showed that 85 (86%) were resistant to itraconazole (ITZ), the first-line clinical drug for chronic infections, 63 (64%) were resistant to voriconazole (VCZ), the first-line agent for invasive infections, 18 (18%) were resistant to posaconazole (PCZ) and 82 (83%) were resistant to isavuconazole (ISZ) (**Figure 1D**). Of note, 50 (51%) of AR*Af* were resistant to three medical azoles and 14 (14%) were resistant to all tested medical azoles (**Table S1**), identifying a UK-wide aerosolized exposure to drug-resistant variants of this pathogen.

**Figure 1:**
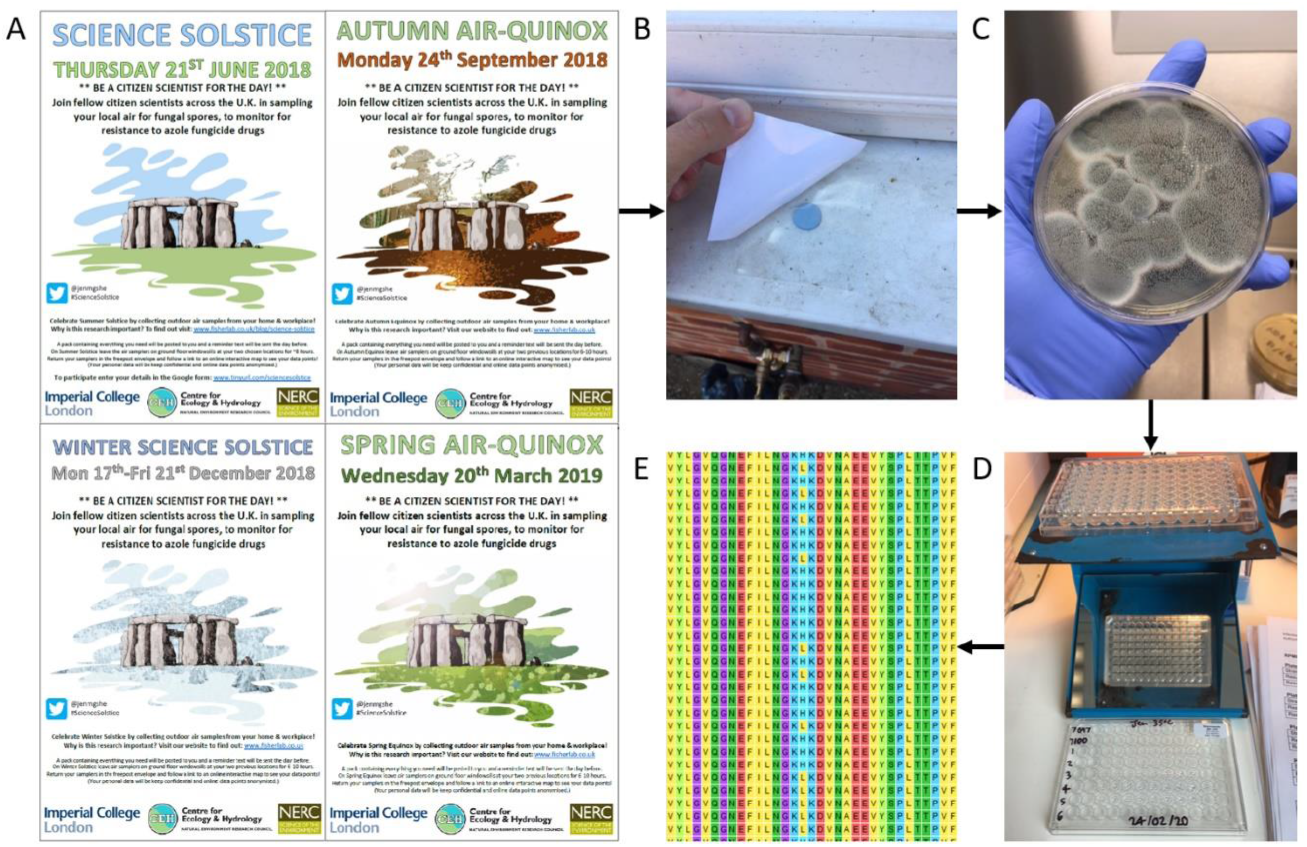
A) Posters advertising the four air sampling rounds that were posted on social media and displayed at Imperial College London and UK CEH, B) photo of a passive air sampler attached to an outdoor ground floor windowsill that had its sticky side exposed for 6-10 hours on a sampling day, C) isolates of *Aspergillus fumigatus* grown from an air sampler following incubation at 43°C for 48 hours, D) minimum inhibitory concentration (MIC) testing of tebuconazole-resistant *A. fumigatus* isolates to medical azoles, E) amino acid sequences spanning the L98H substitution for a subset of tebuconazole-resistant *A. fumigatus* isolates.

**Figure 2.**
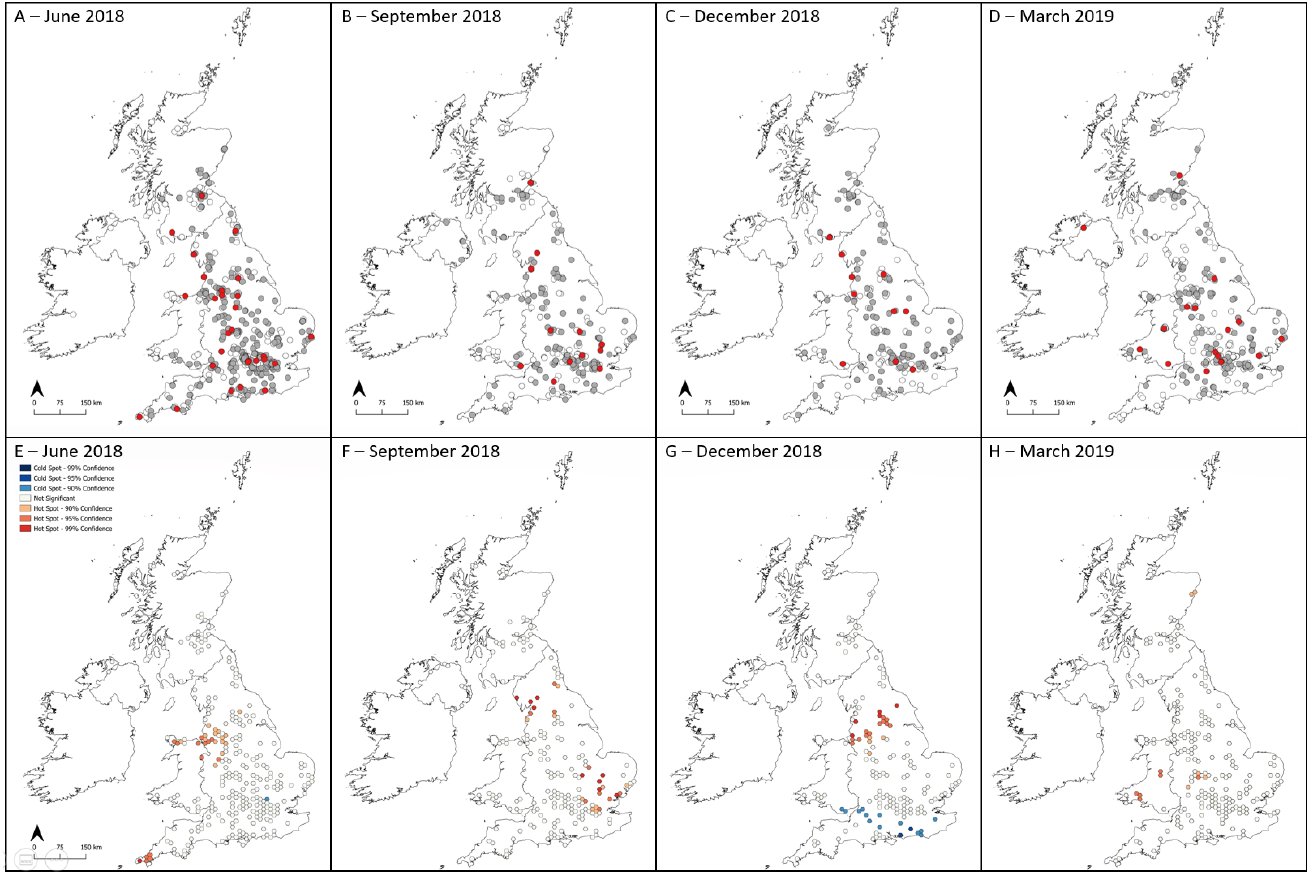
Maps showing locations of UK and Republic of Ireland air samples collected on A) 21^st^ June equinox 2018, B) 24^th^ September solstice 2018, C) 21^st^ December equinox 2018 and D) 20^th^ March solstice 2019. White dots indicate samples that did not grow *A. fumigatus*, grey dots indicate samples that grew tebuconazole-susceptible *A. fumigatus* and red dots indicate samples that grew tebuconazole-resistant *A. fumigatus*. Hotspots of azole-resistant *A. fumigatus* with 90%, 95% and 99% confidence according to Getis-Ord Gi* cluster detection analysis for sampling rounds on E) 21^st^ June 2018, F) 24^th^ September 2018, G) 21^st^ December 2018 and H) 20^th^ March 2019.

**Table 1:**
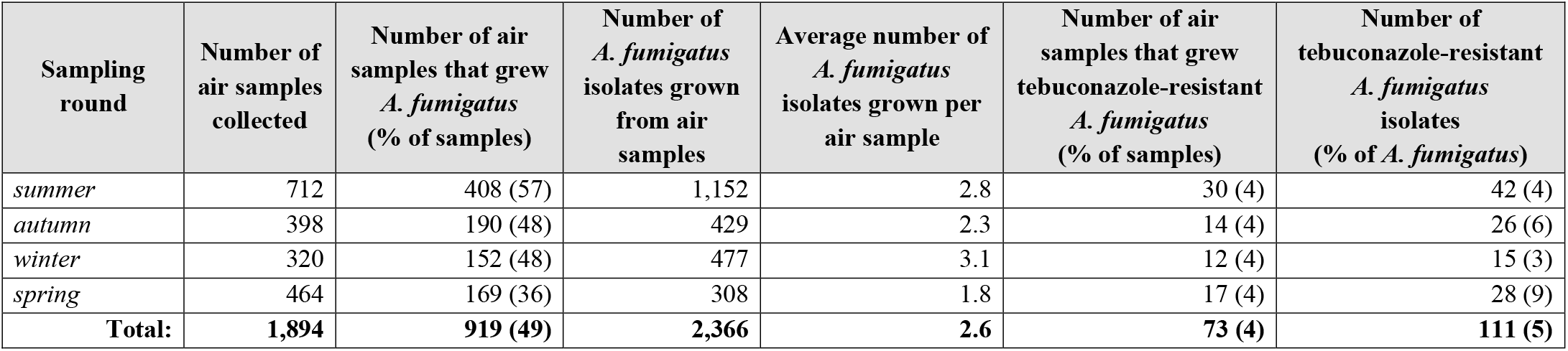
The number of air samples collected, the number of air samples that grew *A. fumigatus* and the number of samples that grew azole-resistant *A. fumigatus* across the four air sampling rounds.

In order to examine the genetic basis of the TEB-resistant phenotype, we genotyped the canonical locus that confers azole-resistance, the sterol-demethylase gene *cyp51A* (*19, 20*). Environmental azole resistance in *A. fumigatus* is most commonly due to within-gene point mutations in *cyp51A* (**Figure 1E**) that are twinned with expression-upregulating tandem repeats (TRs) in the promoter region (*21*). Genotyping confirmed that the most common aerosolized resistance-associated polymorphisms were TR_34_/L98H (59%) and TR46/Y121F/T289A (6%). We further found that 30% of TEB-resistant isolates did not contain any polymorphisms in the *cyp51A* promoter or coding regions suggesting the existence of alternative resistance mechanisms as have previously been noted (*22*) (**Table S1**). This is of concern in the clinical setting as only the two former mutations are picked up by commercially available PCR diagnostic methods, although previously unidentified mutations leading to resistance will be picked up by phenotypic testing of minimum inhibitory concentration (MIC) providing that there is an isolate available to test.

Subsequently, we determined the extent to which aerosolized AR*Af* matched those previously recovered and sequenced from the UK terrestrial environments and patient cohorts by sequencing the genomes of 62 AR*Af* isolates from the summer 2018 air sampling round (21 wild-type; 41 TEB-resistant). These data were then combined with those resulting from prior UK-wide genomic surveillance (*23*) in a phylogenetic analysis. The resulting tree showed that aerosolized isolates were broadly distributed throughout the UK phylogeny (**Figure 3A**) and were drawn from both previously described clades ‘*A*’ (which contains the majority of AR*Af* (*23*)) and ‘*B*’ (which is mainly sensitive to azole fungicides). Principal component analysis (**Figure 3B**) corroborated our conclusion that there was no clear differentiation between aerosolized isolates when compared to those from clinical and terrestrial sources, and nucleotide diversity (*π*) tests showed that the observed genetic diversity separating these groups was not significantly different (one-tailed t-test *p* < 0.12591). We did, however, note two clusters of aerosolized isolates that were largely unrepresented in our previous surveillance, suggesting a temporally dynamic aspect to the UK population genetic structure of this fungus (**Figure 3A**). When mapped against the *A. fumigatus* reference genome *Af293*, each pair of isolates across the combined dataset were, on average, separated by 24,000 SNPs, a figure that was marginally lower (23,250 SNPs) when considering only aerosolized isolates. Significantly, the genotype of an aerially sourced AR*Af* isolate grouped within a previously identified UK-wide clonal subclade of genotypes, Clade A*_A_*. Isolates in this clade all bear the hallmark TR_34_/L98H resistance allele and are widely found in the environment and infecting patients. From these genomic data, we concluded that the genotypes of AR*Af* recovered from the UK aerobiome are largely (but not exclusively) representative of those azole-resistant genotypes recovered from patients. Moreover, these data indicate that ~40% (58/150 sequenced genomes) of AR*Af* infections are acquired from environmental exposures.

**Figure 3.**
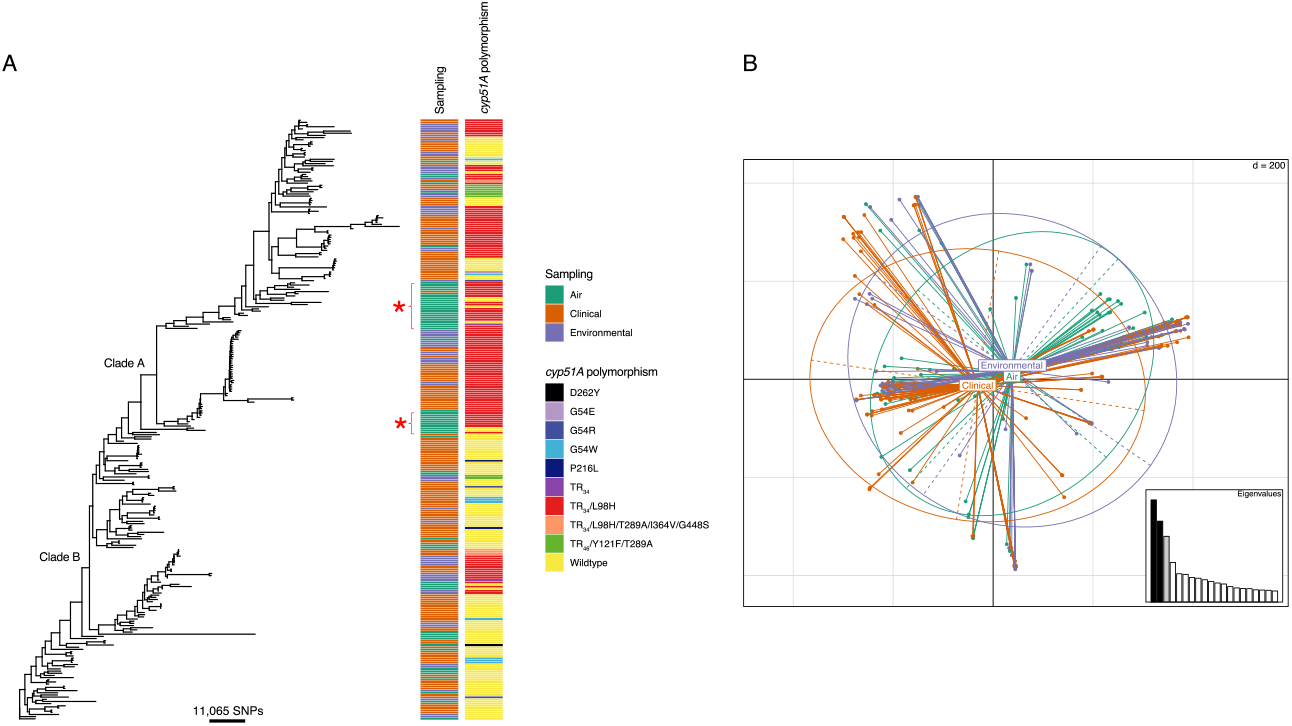
Phylogenetic and population genetic analyses indicate isolates sampled from air are not genetically distinct from the wider *Aspergillus fumigatus* population. A) Unrooted maximum likelihood phylogenetic tree (constructed in RAxML using genome-wide SNPs) showing the sampling type and *cyp51a* polymorphisms. Isolates sampled from air are found throughout the phylogeny. B) PCA indicate genetic identity of isolates, with air sampled isolates broadly deriving from the same wider *A. fumigatus* population.

We next sought to explain the spatial occurrence of *A. fumigatus* and AR*Af* using logistic regression to determine which cardinal environmental variables (season, maximum daily temperature, land cover classification and proximity to nearest industrial composting facility with open windrow or outdoor activity (OW/OA)) affected their likelihood. Individually, each variable was found to have a significant effect on whether a sample grew the mold (season *χ*^2^ = 50.3, df = 3, *p* < 0.01; maximum daily temperature *χ*^2^ = 29.3, df = 1, *p* < 0.01; land cover classification *χ*^2^ = 17.4, df = 7, *p* = 0.01; proximity to nearest OW/OA composting facility *χ*^2^ = 15.1, df = 1, *p* < 0.01; **Table 2**). Negative binomial regression then determined which environmental variables affected the number of *A. fumigatus* colonies grown, with sampling round (*χ*^2^ = 35.9, df = 3, *p* < 0.01) and proximity of sampling location to the nearest OW/OA composting facility (*χ*^2^ = 10.2, df = 1,*p* < 0.01) showing significant associations. Subsequently, Average Nearest Neighbour (ANN) tests found sampling locations in each sampling round to be clustered (**Table 3**) and Getis-Ord Gi* spatial clustering analysis detected hotspots (high-) and coldspots (low-prevalence) of airborne AR*Af* in each sampling round (**Figure 2E–H**). However, none of the included environmental variables were found to affect the number of AR*Af* grown from the air-samples, and the detected clusters were not stable between sampling rounds.

**Table 2:** Odds ratios, confidence intervals and p-values calculated for independent variables included in a logistic regression model using air samples collected in the UK (*n* = 1,894) to explain whether a sample grew *A. fumigatus*. Significant results (*p* <= 0.05) are highlighted in bold. OW/OA = open windrow or outdoor activity.

| Independent variable | Odds ratio (95% CI) | Pr(>\|z\|) |
| --- | --- | --- |
| Sampling round |  |  |
| *summer (baseline)* |  |  |
| *autumn* | **0.75 (0.57-0.99)** | **0.05** |
| *winter* | 0.94 (0.57-1.57) | 0.82 |
| *spring* | **0.53 (0.37-0.75)** | **<0.01** |
| Maximum daily temperature at sampling location on sampling date | 1.04 (0.99-1.09) | 0.13 |
| Proximity of sampling location to nearest OW/OA composting facility | **0.99 (0.99-0.99)** | **<0.01** |

**Table 3:** Results for Average Nearest Neighbour tests run for each sampling round separately and all sampling rounds together.

|  | Sampling round |  |  |  |  |
| --- | --- | --- | --- | --- | --- |
|  | summer | autumn | winter | spring | all |
| *Number of samples* | 712 | 398 | 320 | 463 | 1893 |
| *Observed Mean Distance (m):* | 3,059 | 3,505 | 3,314 | 2,945 | 855 |
| *Expected Mean Distance (m):* | 10,544 | 14,103 | 15,728 | 13,075 | 6,467 |
| *Nearest Neighbour Ratio:* | 0.29 | 0.25 | 0.21 | 0.23 | 0.13 |
| *z-score:* | -36.24 | -28.68 | -27.01 | -31.89 | -72.23 |
| *p-value:* | <0.00 | <0.00 | <0.00 | <0.00 | <0.00 |
| *Pattern* | Clustered | Clustered | Clustered | Clustered | Clustered |

In parallel to our aerosol sampling, a soil-sampling campaign was conducted by citizen scientists during the 2019 summer solstice, resulting in the recovery of a high burden of AR*Af* from garden soils totaling 736 (14%) resistant isolates from 246 locations (*8*). There were 46 participants from which both soil and at least one air sample were collected (**Supplementary Table 2**). Of these, 23 (50%) grew AR*Af* colonies from either an air sample or soil sample, but only 3 (7%) grew AR*Af* colonies from both. Moreover, there were no locations from which AR*Af* was isolated from an air sample more than once. Taken together, these observations lead us to conclude that the locally dynamic nature of atmospheric flows means that our bioaerosol sampling strategy does not capture the presence of local AR*Af* soil hotspots. The corollary of this observation is that atmospheric mixing leads to the UK population being, on average, equally exposed to this bioaerosol. Based on our air sampling data and this assumption, we estimate that the cumulative exposure of each individual across the UK to AR*Af* averages 22 (95% CI: 6-38) days per year.

Importantly, while our study only reports on aerosolized spores collected in the UK, this exposure is not restricted to the UK. Recruitment of citizen scientists attracted several participants from outside the UK (*24*) with air samples from Germany, France and The Netherlands growing viable AR*Af*. Fungal spores are readily dispersed across intercontinental scales (*25*), and our population genetic studies of *A. fumigatus* has shown that its worldwide distribution is unstructured, with no evidence for isolation-by-distance effects (*26*). Moreover, the usage of DMIs in agriculture is widely increasing where measured (e.g., ((*27*)), and environmental azole-resistance has been detected in every country in which it was monitored (*28*) indicating that AR*Af* has achieved a global occurrence. Yet, to date, there has been no attempt to systematically measure population-wide exposures to AR*Af* more globally. This is changing, and in 2022 the pan-South American LatAsp (https://www.latasp.com) surveillance study commenced, mirroring the essential features of our Citizen Science campaign, with the aim of determining the continent-wide incidence of AR*Af*. Nonetheless, identifying the ecological hotspots that ultimately cause the miasma of azole-adapted mould that we document will require spatially downscaled approaches if they are to be mitigated.

## Methods

### *Culturing* Aspergillus fumigatus *from UK air samples*

Air samples from which *A. fumigatus* isolates were cultured for this study were collected in four sampling rounds of a citizen science project that took place between June 2018 and March 2019. The four sampling rounds took place on 21^st^ June 2018 (summer solstice), 24^th^ September 2018 (autumn equinox), 21^st^ December 2018 (winter solstice) and 20^th^ March 2019 (spring equinox). In total, 485 individuals collected 1,894 air samples from England, Wales, Scotland and Northern Ireland; 712 samples in summer, 398 samples in autumn, 320 samples in winter and 464 samples in spring (**Table 1**).

Prior to the sampling date, citizen scientists were posted two passive air samplers (measuring 6.8 cm x 8.0 cm) to collect *A. fumigatus* spores, which were MicroAmp^™^ clear adhesive films (ThermoFisher Scientific, UK) cut in half. The sticky side of each sampler was exposed horizontally for 6-10 hours at approx. 1 m height, on the sampling date, re-covered and returned by post to the primary author. When an air sample was received it was stored at room temperature until processing, which involved removing the cover and placing the sampler sticky-side down on a Sabouraud dextrose agar (SDA; Merck, Germany) plate containing penicillin (Merck, Germany) at 200 mg/L and streptomycin (Merck, Germany) at 400 mg/L. The plate was incubated at 43°C for 24 hours, the sampler was removed, and the plate was incubated for a further 24 hours at 43°C. *A. fumigatus* isolates were picked one at a time using a sterilized wooden toothpick into a tube containing mold preservation solution (MPS; 0.2% agar and 0.05% Tween 20 in dH_2_O) and stored at 4°C.

### Isolate screening for azole resistance

Isolates were screened for tebuconazole resistance by pipetting 5 μl of MPS containing *A. fumigatus* spores onto an SDA plate containing 6 mg/L tebuconazole, and a subset of isolates (*n* = 250) were also tested using the Tebucheck protocol (*17*). The concentration of 6 mg/L tebuconazole was chosen after testing the growth of 30 isolates with known *cyp51A* mutations on SDA supplemented with 0 mg/L, 4 mg/L, 6 mg/L, 8 mg/L and 16 mg/L tebuconazole. Isolates able to grow at a tebuconazole concentration of 6 mg/L were tested for susceptibility to ITZ, VCZ, PCZ and ISZ according to CLSI M38-A2, as described in Borman *et al*. (2017). Minimum inhibitory concentrations (MICs) were recorded as the lowest drug concentration at which no growth was observed, and MICs were considered resistant when they were >1 mg/L for ITZ, VCZ and ISZ and >0.25 mg/L for PCZ which are the suggested clinical breakpoints.

### *Identification of* A. fumigatus cyp51A *gene azole-resistance alleles*

The promoter region of *cyp51A* was amplified using forward primer 5’-GGACTGGCTGATCAAACTATGC-3’ and reverse primer 5’-GTTCTGTTCGGTTCCAAAGCC-3’ and the PCR conditions: 95°C for five minutes; 30 cycles of 98°C for 20 seconds, 65°C for 30 seconds and 72°C for 30 seconds; 72°C for five minutes. The PCR reaction volume used was 50 μl: 10 μl of FIREPol^®^ DNA polymerase (Solis Biodyne, Estonia), 10 μl of forward primer (1.5 μM; Invitrogen, US), 10 μl of reverse primer (1.5 μM; Invitrogen, US), 18 μl of nuclease-free water (Merck, Germany) and 2 μl of DNA. Amplicons were visualized by gel electrophoresis and samples with visible bands were sent for sequencing using the forward primer. The coding region of *cyp51A* was amplified using forward primer 5’-ATGGTGCCGATGCTATGG-3’ and reverse primer 5’-CTGTCTCACTTGGATGTG-3’ and the PCR conditions: 94°C for two minutes; 35 cycles of 94°C for 30 seconds, 60°C for 45 seconds and 72°C for 45 seconds; 72°C for five minutes. The PCR reaction volume used was 50 μl: 0.2 μl of Q5® high-fidelity DNA polymerase (New England Biolabs, UK), 10 μl of Q5® reaction buffer (5X; New England Biolabs, UK), 0.5 μl of deoxynucleotide (dNTP) solution mix (40 μM; New England Biolabs, UK), 1 μl of forward primer (10 μM; Invitrogen, US), 1 μl of reverse primer (10 μM; Invitrogen, US), 35.3 μl of nuclease-free water (Merck, Germany) and 2 μl of DNA. Amplicons were visualized by gel electrophoresis and samples with visible bands were sent for sequencing in two segments using the primers 5’-CTGATTGATGATGTCAACGTA-3’ and 5’-GATTCACCGAACTTTCAAGGCTCG-3’ (*29*). Sequences were aligned using Molecular Evolutionary Genetics Analysis (MEGA) software (Penn State University, US).

### Identification of isolates

Isolates that failed to sequence using the primers for the promoter and coding regions of *cyp51A* were identified using matrix assisted laser desorption ionization-time of flight (MALDI-TOF) mass spectrometry (MS), as described by Fraser and colleagues. (2016).

### *Whole-genome sequencing of* A. fumigatus *isolates*

Genomic DNA (gDNA) was extracted from tebuconazole-resistant isolates with identity confirmed as *A. fumigatus*. Isolates were revived from cryopreservation at −80°C by pipetting 20 μl into 25 cm^3^ Nunc^™^ flasks containing SDA and incubating at 37°C for 48 hours. Spores were harvested by washing the surface of the SDA with 10 ml of phosphate-buffered saline (PBS) plus 0.01% Tween 20, 1.8 ml of spore suspension was added to 2 ml FastPrep tubes (MP Biomedicals, US) and tubes were centrifuged at 5,000 rpm for 10 minutes. The supernatant was discarded and the pellet resuspended in 300 μl of lysis solution and 1 μl of RNase A from the MasterPure^™^ Complete DNA and RNA Purification Kit (Lucigen, US). The kit protocol was followed, including an additional bead-beading step using a FastPrep-24^™^ instrument. Extracted gDNA was purified using a DNeasy Blood and Tissue Kit (Qiagen, Germany) and DNA concentration was measured using a Qubit fluorometer and Qubit dsDNA BR Assay kit (ThermoFisher Scientific, UK). A NanoDrop^™^ spectrophotometer (ThermoFisher Scientific, UK) was used to assess DNA purity by checking that the ratio of absorbances at 260/230 nm and 260/280 nm were 1.8-2.0. Purified gDNAs were stored at −20°C prior to being sent to Earlham Institute (UK) where gDNA libraries were constructed, normalised and indexed. Libraries were run on a NovaSeq 6000 SP v1.5 flow cell to generate 150 bp paired-end reads. These data are deposited in the European Nucleotide Archive (ENA) under Project Accession PRJEB51237.

All raw reads were quality checked with FastQC v0.11.5 (Babraham Institute) and aligned to the reference genome Af293 (*30*) using Burrows-Wheeler Aligner v0.7.8 MEM (*31*) before conversion to sorted BAM format using SAMtools v1.3.1 (*32*). Variant calling was performed with GATK HaplotypeCaller v4.2.6.1 (*33*), excluding repetitive regions (identified by RepeatMasker v4.0.6), generating GVCFs. Low-confidence variants were filtered providing they met at least 1 of the parameters DP < 5, GQ < 50, MQ < 40, MQRankSum < −12.5, ReadPosRankSum < −8.0, SOR > 4.0. In addition, alternate variants must be present in at least 90% of the reads. SNPs were mapped to genes using vcf-annotator (Broad Institute).

Phylogenetic analysis was carried out on 62 tebuconazole-resistant *A. fumigatus* isolates collected by air sampling in addition to 215 environmental and clinical *A. fumigatus* isolates with complete sampling history collected in the UK between 2005 and 2017 (*23*) (data available from PRJEB27135 and PRJEB8623). Whole genome SNP data were converted to presence/absence of a SNP with respect to reference, and any SNPs identified as low confidence in the variant filtration step were assigned as missing. These data were converted to FASTA format. Maximum-likelihood phylogenies were constructed using rapid bootstrap analysis over 1000 replicates using the GTRCAT model of rate heterogeneity in RAxML v8.2.9 (*34*) to assess sequence similarity between isolates, and resulting phylogenies were visualized using ggtree v3.1.14 and Microreact (https://microreact.org/project/6NMrDobYGZnhmYnMSsHhC5-air) (**Figure S1**).

Genetic similarities were investigated using the hypothesis-free approaches Principal Component Analysis (PCA) and Discriminant Analysis of Principal Components (DAPC) (*35*) using the R package adegenet v2.1.5 in R v4.1.0. Nucleotide diversity tests were implemented in VCFtools v0.1.13 (*36*).

### *Environmental variables thought to influence growth of* A. fumigatus

**Table 4** details the environmental variables that were ascertained for the sampling dates and locations, based on the information provided by citizen scientists. Sampling round was used as a proxy for season: 21^st^ June 2018 (summer), 24^th^ September 2018 (autumn), 21^st^ December 2018 (winter), 20^th^ March 2019 (spring).

**Table 4:**
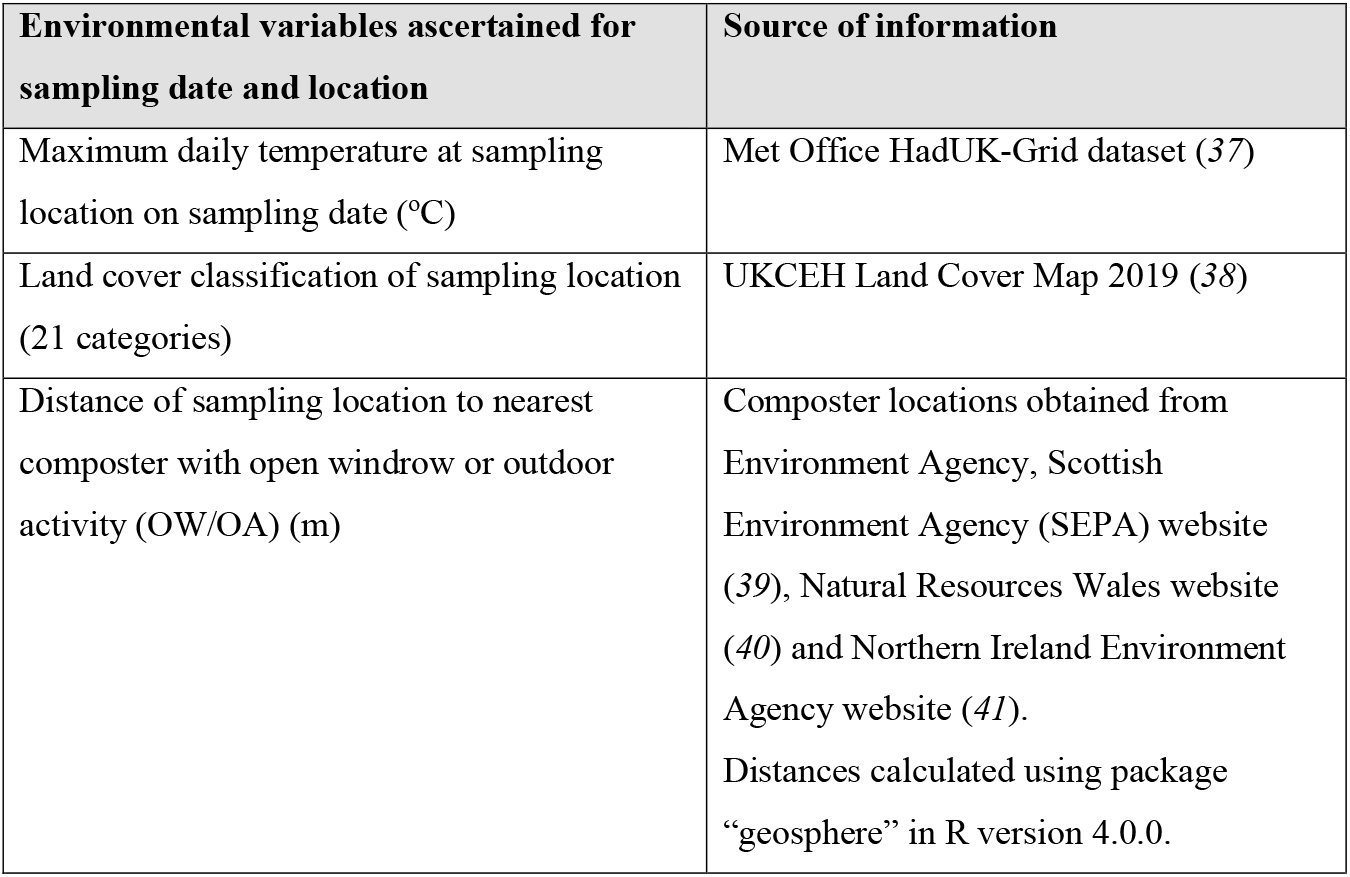
Environmental variables obtained for air sampling locations and dates and the sources they were obtained from.

### Generalized linear models

Generalized linear models (GLMs) run in R v4.0.0 (*42*) were used to find associations between the environmental variables in **Table 4** and i) the likelihood of a sample growing *A. fumigatus*, and ii) likelihood of a sample growing AR*Af*. Growth of *A. fumigatus* or AR*Af* from a sample was categorized as 0/1 and logistic regressions (“glm” function; family = “binomial”) were performed. Significant improvement on the null model, as determined by analysis of variance (ANOVA) using chi-squared test, determined which environmental variables were included in the regression model. Reduced Akaike information criterion (AIC) score and significant improvement on the null model were used to choose the regression model with the best fit. Results were considered significant when p <= 0.05.

### Spatial clustering analysis

ArcMap 10.7 was used to investigate the spatial distribution of all sampling locations and the samples that grew AR*Af*. Sample locations were georeferenced and projected using the British National Grid (EPSG:27700) coordinate reference system.

The Average Nearest Neighbour (ANN) test was used to determine whether the sampling locations in each sampling round, and across all sampling rounds, were randomly distributed or spatially autocorrelated. This test measures the distance between each sampling location and the next closest sampling location and compares this to the distance that would be expected if the locations were randomly distributed throughout the study area (the UK). If the observed mean distance is less than the expected mean distance, the spatial distribution of the observed data is considered spatially clustered. Alternatively, if the mean distance is greater than the expected distribution, the features are considered dispersed. In both scenarios, ANN ratios were used to establish the type of spatial pattern (*i.e*. clustered, dispersed or random) and associated p values <0.05 were used to establish statistical significance.

A local indicator of spatial autocorrelation was used to detect and geographically identify hotspots of azole resistance in the UK. First, sampling locations were geospatially aggregated within a gridded hexagon structure (cell area 115km^2^). For each cell, the total number of collected samples and the total number of azole-resistant *A. fumigatus* samples were extracted and used to calculate sample positivity (%). Cells with zero sample locations were removed prior to analysis. Then, a Getis Ord Gi* analysis was conducted to identify statistically significant hotspot cells based on sample positivity. This approach was used to generate local z scores for each cell, with statistically significant high and low values indicating hotspots and coldspots, respectively. For this analysis hot- and coldspot cells were generated and grouped using three different significance values (90, 95 and 99%).

### Average exposure simulation analysis

Assuming that any location could potentially be AR*Af* positive at some point in time, we conducted a simulation based on 1,000,000 individuals, each having a respiratory rate extracted from a normal distribution with a mean of 9L/min and a standard deviation of 2L/min. To match the air sampling campaigns conducted, we considered 1,095 time periods of 8 hours over a period of one year (365 days). The probability of each individual being exposed to AR*Af* was based on a truncated normal distribution of mean 5.86% and standard deviation of 2.80%, set with 0 as the lower bound. We then calculated the number of 8-hour AR*Af* positive period for each individual, from which we derived the annual average across our total population.

## Acknowledgements

The authors would like to thank all the citizen scientists who collected air samples for this study. We also thank Dr Pippa Douglas for providing the locations of composters in England with open windrow or outdoor activity, and Dr Jianhua Zhang for sharing the *cyp51A* coding region primers.

## Author Contributions

JMGS, ACS, and MCF conceptualised the study; AA, PSD, MF, AMB and EMJ contributed experimental techniques; JMGS, SH, APB and TRS processed samples; JMGS, JR, CBU and FBP analysed the data; JMGS, JR and MCF drafted the original manuscript, which ACS, EMJ, SH and FBP reviewed and edited.

