## Supplemental Files for "Landscape-scale exposure to multiazole-resistant *Aspergillus fumigatus* bioaerosols"

**Figure S1.** Sampling locations and maximum-likelihood phylogenies constructed from whole-genome data of 62 air-sampled *Aspergillus fumigatus* isolates and 215 environmental and clinical *A. fumigatus* isolates collected in the UK between 2005 and 2017 visualized using Microreact (https://microreact.org/project/6NMrDobYGZnhmYnMSsHhC5-air).

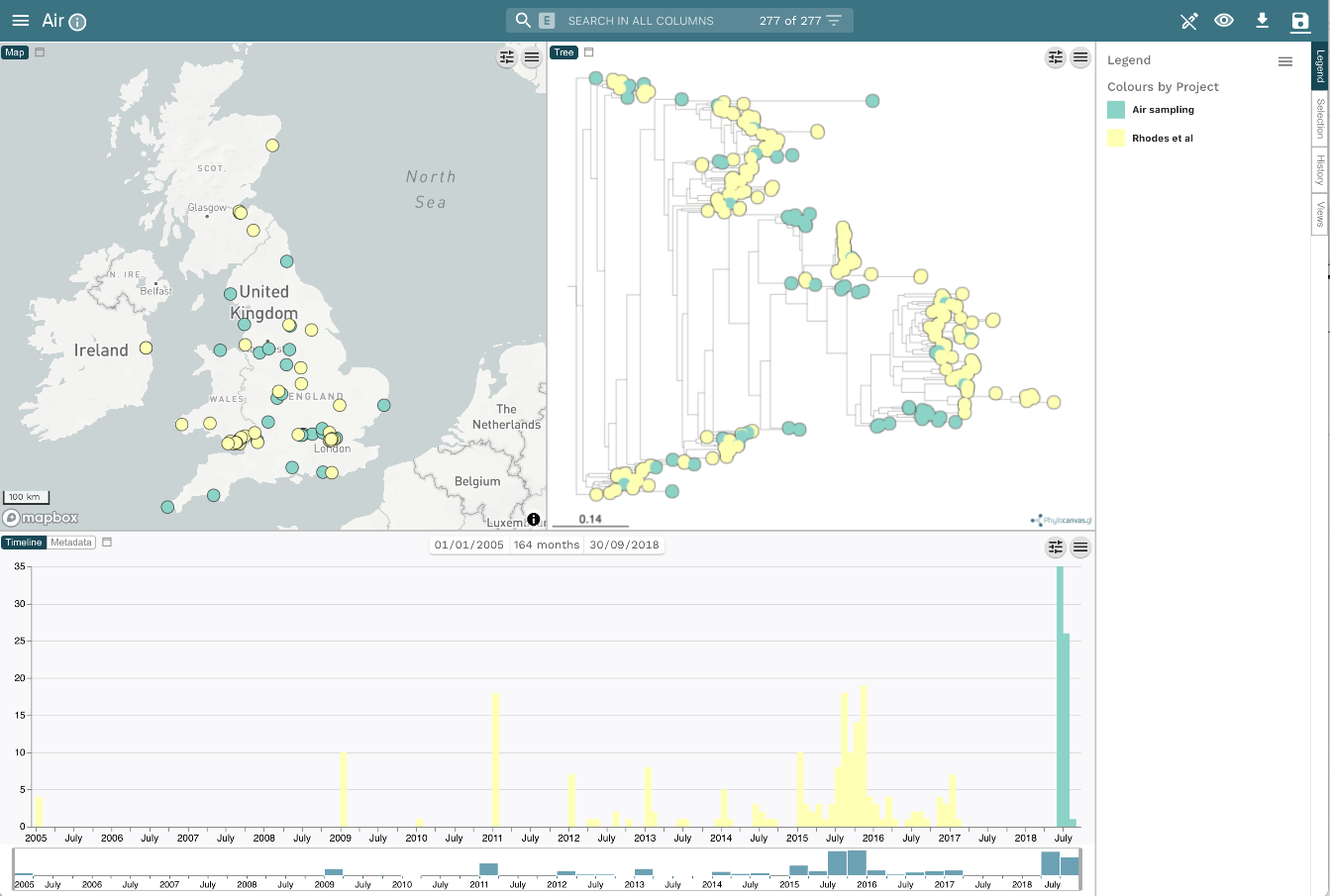

| **Supplementary Table 1:** Details of the 111 UK tebuconazole-resistant isolates including the unique identifier for the air sample they were grown from, the county from which the sample was collected, their minimum inhibitory concentrations (MICs) for itraconazole (ITZ), voriconazole (VCZ), posaconazole (PCZ) and isavuconazole (ISZ), Tebucheck result and *cyp51A* polymorphisms. Sample prefix indicates air sampling round that sample was collected in: SS18=21^st^ June 2018, AE18=24^th^ September 2018, WS18=21^st^ December 2018, SA19=20^th^ March 2019. Resistant MICs are highlighted in bold and isolates subsequently found not to be *Aspergillus fumigatus* are highlighted in red. WT=wild-type. ND=not done. | | | | | | | |
| --- | --- | --- | --- | --- | --- | --- | --- |
| **Sample**  **(prefix for sampling round)** | **County,**  **north to south** | **ITZ** | **VCZ** | **PCZ** | **ISZ** | **TebuCheck result (mg/L)** | ***cyp51A* polymorphisms** |
| SS18-084-02 | Edinburgh | **16** | **2** | 0.25 | **4** | 16 | TR_34_/L98H |
| SS18-251-01 | Northumberland | **16** | **2** | 0.25 | **2** | - | TR_34_/L98H |
| SS18-073-01 | Dumfries & Galloway | 0.25 | **2** | 0.12 | 0.5 | 16 | *A. lentulus* |
| SS18-074-02 | Cumbria | **16** | **2** | **0.5** | **2** | 16 | TR_34_/L98H |
| SS18-068-01 | Lancashire | **16**  **16** | **2**  **2** | 0.25  0.25 | **2**  **2** | 8  6 | TR_34_/L98H  TR_34_/L98H |
| SS18-007-02 | West Yorkshire | **16** | **2** | 0.25 | **2** | 16 | TR_34_/L98H |
| SS18-381-02 | Cheshire | **16** | **2** | 0.25 | **2** | - | TR_34_/L98H/N479S |
| SS18-202-02 | South Yorkshire | **16** | **4** | 0.25 | **2** | - | TR_34_/L98H |
| SS18-403-01 | Conway | **16** | **2** | 0.25 | **2** | - | WT |
| SS18-075-01 | Cheshire | **16** | **4** | **0.5** | **4** | 8 | TR_34_/L98H |
| SS18-257-01 | Derbyshire | **16** | **2** | **0.5** | **2** | - | TR_34_/L98H |
| SS18-261-02 | West Midlands | **16**  0.12 | **2**  0.5 | **0.5**  0.06 | **2**  0.5 | -  - | TR_34_/L98H  WT |
| SS18-278-02 | West Midlands | **16** | 1 | **0.5** | **2** | - | TR_34_/L98H |
| SS18-085-01 | Worcestershire | **16** | **2** | 0.25 | **4** | 16 | TR_34_/L98H |
| SS18-139-01 | Suffolk | **16** | **2** | 0.25 | **4** | - | TR_34_/L98H |
| SS18-177-02 | Gloucestershire | **16** | **2** | 0.25 | **2** | - | WT |
| SS18-143-01 | Hertfordshire | **16** | **2** | 0.25 | **4** | - | TR_34_/L98H |
| SS18-082-01 | Hertfordshire | **16**  **16**  **16**  **16** | **4**  **4**  **2**  **2** | **0.5**  **0.5**  0.25  0.25 | **4**  **4**  **4**  **2** | 16  8  6  8 | TR_34_/L98H  TR_34_/L98H  TR_34_/L98H  TR_34_/L98H |
| SS18-056-02 | Buckinghamshire | **16**  **16**  **16**  **16** | 1  **2**  **2**  **2** | **0.5**  0.25  0.25  0.25 | **2**  **2**  **2**  **2** | 6  6  8  8 | TR_34_/L98H  TR_34_/L98H  TR_34_/L98H  TR_34_/L98H |
| SS18-005-02 | Oxfordshire | **16** | **16** | **0.5** | **16** | 16 | TR_46_/Y121F/T289A |
| SS18-013-01 | Oxfordshire | **16** | **2** | 0.25 | **2** | 8 | TR_34_/L98H |
| SS18-040-01 | Oxfordshire | **16** | **2** | **0.5** | **2** | 8 | TR_34_/L98H |
| SS18-375-02 | Oxfordshire | 0.25 | 1 | 0.06 | 1 | - | *A. lentulus* |
| SS18-340-01 | London | **16**  **16** | **2**  **2** | 0.25  0.25 | **2**  **2** | -  - | TR_34_  TR_34_/L98H |
| SS18-423-01 | Bristol | 0.12 | 0.12 | 0.03 | 0.25 | - | WT |
| SS18-304-01 | Hampshire | **16** | **2** | 0.25 | **2** | - | TR_34_/L98H |
| SS18-193-02 | West Sussex | **16**  0.12 | 1  0.25 | **0.5**  0.03 | **2**  0.5 | -  - | TR_34_/L98H  WT |
| SS18-248-01 | Hampshire | **16** | **16** | **0.5** | **16** | - | *A. lentulus* |
| SS18-204-01 | Devon | **16**  **16**  **16** | **2**  **2**  **2** | **0.5**  0.25  **0.5** | **2**  **2**  **2** | -  -  - | TR_34_/L98H TR_34_/L98H  TR_34_/L98H |
| SS18-285-01 | Cornwall | **16** | **2** | 0.25 | **4** | - | WT |
| AE18-081-01 | Fife | **16** | **4** | 0.25 | **2** | - | TR_34_/L98H |
| AE18-026-01 | Cumbria | **16** | **4** | **0.5** | **4** | - | TR_34_/L98H/D481N |
| AE18-098-02 | Lancashire | **16**  **16** | 0.5  **2** | **0.5**  0.25 | **4**  **2** | - | WT  WT |
| AE18-098-01 | Lancashire | **16**  **16**  **16**  **16**  **16**  **16**  0.25  **16**  **16**  **16** | **2**  **2**  **2**  **2**  **2**  1  0.25  **2**  **2**  **2** | 0.25  0.25  0.25  0.25  0.25  0.25  0.06  0.25  0.25  0.25 | **4**  **2**  **2**  **2**  **2**  **2**  0.25  **2**  **2**  **2** | -  -  -  -  -  -  -  -  -  - | D262Y  WT  WT  WT  WT  WT  WT  TR_34_/L98H  WT  WT |
| AE18-016-02 | Lancashire | **16** | **2** | 0.25 | **2** | - | TR_34_/L98H |
| AE18-060-01 | West Midlands | **16** | **2** | 0.25 | **2** | - | TR_34_/L98H |
| AE18-125-01 | Northamptonshire | **16** | 1 | 0.25 | **2** | - | TR_34_/L98H |
| AE18-124-01 | Essex | **16** | **2** | **0.5** | **2** | - | TR_34_/L98H |
| AE18-037-01 | Hertfordshire | **16** | **2** | 0.25 | 1 | - | *A. lentulus* |
| AE18-082-02 | Hertfordshire | 0.25 | 0.5 | 0.06 | 0.5 | - | WT |
| AE18-211-01 | Oxfordshire | **16** | 1 | 0.25 | **2** | - | TR_34_/L98H |
| AE18-225-02 | Cardiff | 0.06 | 0.13 | 0.03 | 0.12 | - | *A. nidulans* |
| AE18-064-01 | London | **16** | **2** | 0.25 | **2** | - | TR_34_/L98H |
| AE18-187-01 | Wiltshire | **16**  **16**  0.12 | 1  **2**  0.25 | 0.25  **0.5**  0.03 | **2**  **4**  0.25 | -  -  - | TR_34_/L98H  TR_34_/L98H  WT |
| WS18-061-01 | Dumfries & Galloway | 0.12 | 0.12 | 0.03 | 0.25 | - | *A. lentulus* |
| WS18-061-02 | Dumfries & Galloway | 0.25 | 1 | 0.12 | 1 | - | *A. lentulus* |
| WS18-134-02 | Cumbria | **16**  **16** | 1  **4** | 0.25  **0.5** | **2**  **8** | -  - | TR_34_/L98H  TR_34_/L98H |
| WS18-060-01 | West Yorkshire | ND | ND | ND | ND | - | TR_34_/L98H |
| WS18-028-02 | Lancashire | **16** | 1 | 0.25 | **2** | - | TR_34_/L98H |
| WS18-153-01 | Merseyside | **16**  **16** | 1  1 | 0.25  0.25 | **2**  **2** | -  - | TR_34_/L98H  TR_34_/L98H |
| WS18-119-02 | Nottinghamshire | **16** | **2** | 0.25 | **4** | - | TR_34_/L98H |
| WS18-045-01 | Lincolnshire | **16** | 1 | 0.25 | **2** | - | TR_34_/L98H |
| WS18-045-02 | Lincolnshire | **16** | 1 | 0.25 | **2** | - | TR_34_/L98H |
| WS18-040-01 | Oxfordshire | **16**  0.06 | **2**  0.25 | 0.25  0.03 | **2**  0.5 | -  - | TR_34_/L98H  D262Y |
| WS18-161-01 | Mid Glamorgan | 0.5 | 1 | 0.12 | 1 | - | *A. lentulus* |
| WS18-125-01 | Surrey | **16** | 1 | 0.25 | **2** | - | TR_34_/L98H |
| SA19-182-02 | Dundee | 0.12 | 0.12 | 0.03 | 0.25 | - | WT |
| SA19-178-01 | Londonderry | 0.06 | 0.12 | 0.03 | 0.25 | - | *A. nidulans* |
| SA19-122-01 | West Yorkshire | **16**  **16** | 1  1 | 0.25  0.25 | 1  1 | -  - | *A. lentulus*  *A. lentulus* |
| SA19-184-01 | Cheshire | 0.06 | 0.25 | 0.03 | 0.25 | -  - | S46F/R66W/  M172V/E427K |
| SA19-050-01 | Staffordshire | **16** | 1 | 0.25 | **2** | - | D262Y |
| SA19-051-01 | Cambridgeshire | **16** | 1 | 0.25 | **2** | - | WT |
| SA19-170-01 | Powys | **16** | **16** | 0.25 | **16** | - | TR_46_/Y121F/T289A |
| SA19-207-02 | Northamptonshire | 0.25 | **16** | 0.12 | **16** | - | TR_46_/Y121F/T289A |
| SA19-176-01 | Suffolk | **16**  **16**  **16**  0.06  0.12  0.06  0.06  0.06 | 1  1  **2**  0.25  0.25  0.25  0.25  0.25 | 0.25  0.25  0.25  0.03  0.03  0.03  0.03  0.03 | **2**  **2**  **2**  0.25  0.5  0.25  0.25  0.25 | -  -  -  -  -  -  -  - | WT  TR_34_/L98H  WT  WT  WT  WT  WT  WT |
| SA19-154-02 | Carmarthenshire | **16**  **16** | **16**  1 | 0.25  0.25 | **8**  **2** | -  - | TR_46_/Y121F/T289A  TR_34_/L98H |
| SA19-001-03 | Oxfordshire | **16** | 1 | 0.25 | **2** | - | TR_34_/L98H |
| SA19-208-01 | Oxfordshire | **16** | **2** | 0.25 | **4** | - | TR_34_/L98H/V486I |
| SA19-063-01 | Essex | 0.12  0.5 | 0.25  0.5 | 0.03  0.03 | 0.25  0.5 | -  - | *A. lentulus*  WT |
| SA19-004-01 | Oxfordshire | 0.25 | 0.25 | 0.03 | 0.5 | - | WT |
| SA19-120-01 | Oxfordshire | **16** | **2** | 0.25 | **2** | - | TR_34_/L98H |
| SA19-078-02 | Mid Glamorgan | **16** | **2** | 0.25 | **2** | - | WT |
| SA19-162-01 | Wiltshire | **16**  **16** | **16**  **16** | 0.25  0.25 | **16**  **16** | -  - | TR_46_/Y121F/T289A  TR_46_/Y121F/T289A |

| **Supplementary table 2:** Details of 46 UK locations from which both an air and soil sample were collected, including the county and the number of *A. fumigatus* (A) and azole-resistant *A. fumigatus* (R) colonies grown from each sample. Samples that grew azole-resistant *A. fumigatus* are highlighted in red and locations that grew azole-resistant *A. fumigatus* from both air and soil samples are highlighted in bold. | | | | | | | | | | | | | | | | | | | | |
| --- | --- | --- | --- | --- | --- | --- | --- | --- | --- | --- | --- | --- | --- | --- | --- | --- | --- | --- | --- | --- |
| **County, north to south** | **Air sampling** | | | | | | | | | | | | | | | | **Soil sampling** | | | |
|  | **Summer 2018** | | | | **Autumn 2018** | | | | **Winter**  **2018** | | | | **Spring 2019** | | | | **Summer**  **2019** | | | |
|  | **1** | | **2** | | **1** | | **2** | | **1** | | **2** | | **1** | | **2** | | **1** | | **2** | |
|  | **A** | **R** | **A** | **R** | **A** | **R** | **A** | **R** | **A** | **R** | **A** | **R** | **A** | **R** | **A** | **R** | **A** | **R** | **A** | **R** |
| Inverness-shire |  |  |  |  | 0 | 0 |  |  | 2 | 0 |  |  | 1 | 0 |  |  | 30 | 0 | 5 | 0 |
| Fife | 1 | 0 |  |  | 3 | 0 |  |  | 0 | 0 |  |  | 1 | 0 |  |  | 0 | 0 | 0 | 0 |
| Midlothian | 2 | 0 | 0 | 0 |  |  |  |  |  |  |  |  |  |  |  |  | 0 | 0 | 30 | 3 |
| **Dumfriesshire** | **0** | **0** |  |  | **0** | **0** |  |  | **7** | **1** |  |  | **0** | **0** |  |  | **1** | **1** |  |  |
| Lancashire | 3 | 0 | 5 | 0 |  |  |  |  |  |  |  |  | 0 | 0 |  |  | 5 | 0 | 3 | 3 |
| Lancashire |  |  |  |  |  |  |  |  |  |  |  |  | 0 | 0 |  |  | 50 | 0 | 0 | 0 |
| West Yorkshire |  |  |  |  | 1 | 0 | 0 | 0 | 34 | 1 | 35 | 0 | 0 | 0 | 0 | 0 | 2 | 0 | 0 | 0 |
| Lancashire |  |  |  |  |  |  |  |  | 8 | 0 | 7 | 0 | 0 | 0 | 0 | 0 | 9 | 0 | 12 | 0 |
| Greater Manchester | 1 | 0 |  |  | 0 | 0 |  |  | 0 | 0 |  |  | 0 | 0 |  |  | 4 | 0 | 2 | 0 |
| Greater Manchester |  |  |  |  |  |  |  |  |  |  |  |  | 0 | 0 | 0 | 0 | 11 | 6 |  |  |
| Cheshire |  |  |  |  |  |  |  |  |  |  |  |  | 0 | 0 |  |  | 0 | 0 | 0 | 0 |
| Cheshire |  |  |  |  |  |  |  |  |  |  |  |  | 1 | 0 |  |  | 0 | 0 | 0 | 0 |
| Cheshire | 2 | 0 | 0 | 0 | 2 | 0 | 0 | 0 | 2 | 0 | 1 | 0 | 0 | 0 | 0 | 0 | 22 | 0 | 50 | 17 |
| Staffordshire |  |  |  |  | 0 | 0 |  |  | 1 | 0 |  |  | 1 | 1 |  |  | 0 | 0 | 0 | 0 |
| Norfolk |  |  |  |  |  |  |  |  |  |  |  |  | 1 | 0 |  |  | 3 | 0 | 0 | 0 |
| West Midlands |  |  |  |  | 2 | 0 |  |  | 0 | 0 |  |  | 0 | 0 | 0 | 0 | 0 | 0 | 0 | 0 |
| West Midlands | 7 | 0 | 8 | 2 | 3 | 0 | 2 | 0 | 0 | 0 | 0 | 0 | 0 | 0 | 0 | 0 | 30 | 0 | 0 | 0 |
| West Midlands | 0 | 0 | 1 | 0 |  |  |  |  |  |  |  |  |  |  |  |  | 3 | 0 | 22 | 0 |
| Northamptonshire |  |  |  |  | 1 | 0 | 1 | 0 | 1 | 0 | 0 | 0 | 1 | 0 | 0 | 0 | 0 | 0 | 2 | 0 |
| Gloucestershire |  |  |  |  |  |  |  |  |  |  |  |  | 0 | 0 | 0 | 0 | 30 | 8 | 30 | 3 |
| Hertfordshire |  |  |  |  | 3 | 0 |  |  | 11 | 0 |  |  | 6 | 0 | 1 | 0 | 50 | 1 | 0 | 0 |
| Hertfordshire |  |  |  |  |  |  |  |  |  |  |  |  | 1 | 0 |  |  | 0 | 0 | 6 | 0 |
| Breconshire | 0 | 0 |  |  | 0 | 0 |  |  |  |  |  |  |  |  |  |  | 0 | 0 | 9 | 0 |
| Oxfordshire | 0 | 0 |  |  | 0 | 0 |  |  |  |  |  |  | 0 | 0 |  |  | 30 | 30 | 6 | 0 |
| Buckinghamshire | 0 | 0 | 1 | 0 | 0 | 0 | 0 | 0 | 0 | 0 | 0 | 0 | 0 | 0 | 0 | 0 | 0 | 0 | 50 | 19 |
| Essex |  |  |  |  |  |  |  |  | 0 | 0 |  |  | 0 | 0 |  |  | 0 | 0 | 0 | 0 |
| Buckinghamshire | 2 | 0 |  |  | 1 | 0 |  |  | 0 | 0 |  |  | 0 | 0 |  |  | 3 | 0 | 30 | 3 |
| Oxfordshire | 1 | 0 |  |  |  |  |  |  |  |  |  |  |  |  |  |  | 30 | 1 | 30 | 0 |
| Oxfordshire | 4 | 0 |  |  | 1 | 1 |  |  | 1 | 0 |  |  | 0 | 0 |  |  | 5 | 0 | 0 | 0 |
| **Oxfordshire** | **0** | **0** | **1** | **0** | **3** | **0** | **1** | **0** | **2** | **0** | **1** | **0** | **1** | **1** | **1** | **0** | **25** | **8** | **25** | **2** |
| Oxfordshire | 0 | 0 |  |  | 0 | 0 |  |  | 0 | 0 |  |  | 0 | 0 |  |  | 30 | 2 | 30 | 6 |
| Greater London | 7 | 0 | 5 | 0 | 2 | 0 | 5 | 0 | 3 | 0 | 0 | 0 |  |  |  |  | 2 | 0 | 17 | 0 |
| Greater London | 1 | 0 | 0 | 0 | 1 | 0 |  |  |  |  |  |  | 0 | 0 |  |  | 19 | 0 | 2 | 0 |
| Buckinghamshire |  |  |  |  |  |  |  |  | 0 | 0 | 3 | 0 | 1 | 0 | 1 | 0 | 4 | 0 | 12 | 0 |
| Greater London | 1 | 0 |  |  | 0 | 0 |  |  |  |  |  |  | 4 | 0 |  |  | 16 | 1 | 8 | 0 |
| Greater London | 0 | 0 |  |  |  |  |  |  |  |  |  |  |  |  |  |  | 25 | 9 | 30 | 5 |
| **Greater London** | **2** | **0** | **1** | **0** | **1** | **1** | **0** | **0** | **0** | **0** | **0** | **0** | **0** | **0** | **0** | **0** | **30** | **1** | **30** | **2** |
| Hampshire | 2 | 0 |  |  | 1 | 0 |  |  | 0 | 0 |  |  | 0 | 0 |  |  | 30 | 0 | 4 | 0 |
| Wiltshire | 0 | 0 | 0 | 0 | 0 | 0 | 1 | 0 | 0 | 0 | 1 | 0 | 0 | 0 | 1 | 0 | 30 | 0 | 0 | 0 |
| Somerset |  |  |  |  | 1 | 0 | 1 | 0 |  |  |  |  | 0 | 0 | 1 | 0 | 0 | 0 | 30 | 3 |
| Dorset |  |  |  |  | 0 | 0 |  |  | 0 | 0 |  |  | 0 | 0 |  |  | 0 | 0 | 0 | 0 |
| Hampshire | 5 | 1 |  |  | 1 | 0 |  |  | 1 | 0 |  |  | 3 | 0 |  |  | 0 | 0 | 0 | 0 |
| Dorset | 5 | 0 | 0 | 0 | 0 | 0 | 0 | 0 | 0 | 0 | 0 | 0 | 1 | 0 | 1 | 0 | 12 | 0 | 30 | 1 |
| Cornwall | 0 | 0 |  |  | 2 | 0 |  |  | 2 | 0 |  |  | 0 | 0 |  |  | 3 | 0 | 3 | 0 |
